# A microbiome assessment of healthy and bleached *Halymenia floresii* (Clemente) C.Agardh (Rhodophyta, Florideophyceae)

**DOI:** 10.64898/2026.02.11.705022

**Authors:** Shareen A Abdul Malik, Santiago Cadena, Abril M. Gamboa-Muñoz, José Q. García-Maldonado, Nathalie Bourgougnon, Daniel Robledo

## Abstract

Macroalgae serve as critical habitat-formers and primary producers in coastal ecosystems, functioning across near-subtidal and intertidal zones in three distinct states: substrate-attached, free-floating (drift), and beach-cast. While substrate-attached macroalgae are susceptible to infectious diseases with significant ecological implications, diseases affecting drift macroalgal communities remain virtually unstudied. Here, we investigated bleaching disease - one of the most common macroalgal afflictions - in the drift rhodophyte *Halymenia floresii* from the Gulf of Mexico. Using 16S rRNA gene high-throughput sequencing and scanning electron microscopy, we characterized the bacterial community structure and composition associated with the free-floating healthy, bleached and degrading *H. floresii* to understand how bacterial partners respond to host health status. Principal Coordinate Analysis based on UniFrac distance revealed distinct clustering of bacterial communities according to host health condition. Shannon diversity indices showed distinct patterns ranging from 1.14 – 3.15 for healthy, bleached, and degrading samples, while Simpson indices ranged from 0.62 to 0.91, reflecting substantial variation in community evenness. In healthy samples, Cyanobacteria (17 – 52%) and Pseudomonadota (previously, Proteobacteria) (41 – 81%) dominated, and the bleached samples were characterized by elevated Bacteroidota (formerly, Bacteroidetes) (5 – 35%) and Pseudomonadota (41 – 88%). Notably, *Novosphingobium* (25 – 49%) dominated healthy hosts while showing lower abundance in degrading (13 – 17%) and bleached (18 – 22%) specimens. Conversely, *Reinekea* emerged as a dominant genus (22.5%) specifically in bleached samples, suggesting a potential role in disease pathogenesis. Microbial network analysis using NetCoMi revealed three distinct bacterial clusters corresponding to health states: a healthy-associated cluster dominated by *Novosphingobium* and uncultured Cyanobacterial with predominantly positive associations, and two disease-associated clusters enriched in opportunistic genera including *Reinekea*, *Vibrio*, *Colwellia*, and *Alteromonas*, indicating network reorganization from cooperative to exploitative interactions. This study provides the first descriptive assessment of microbiome transitions associated with bleaching disease in a drift macroalga and highlights the importance of considering free-floating macroalgal diseases and their potential impacts on coastal ecosystem health.

## 1. Introduction

Macroalgae constitute a dominant component of coastal ecosystem as habitat-formers and primary producers across the near-subtidal and intertidal zones (Longford et al., 2019). Unlike vascular plants, they primarily obtain nutrients from surrounding water and do not require rooting (Lobban and Harrison, 1994; Biber P.D., 2007). While many species remain substrate-attached via holdfasts, natural hydrodynamic forces frequently detach them from their parent populations, forming free-floating drift-mats that eventually become beach-cast wrack or sink to the seafloor (Arroyo and Bonsdroff, 2016). Based on their attachment status, macroalgae function as three distinct ecological groups: (i) substrate-attached populations; (ii) free-floating or drift communities and (iii) beach-cast wrack (Ulaski et al., 2023).

Each functional groups plays distinct ecological roles. Substrate-attached macroalgae form benthic community foundation while drift macroalgae provide additional food resources and habitat complexity in shallow coastal areas (Orth et al., 1984; Pratama et al., 2015; Longford et al., 2019; Hall et al., 2022; Correia et al., 2022; Setia, 2023). The dynamics of drift macroalgal blooms are governed by complex physical, and biological factors, exhibiting distinct seasonal patterns that vary geographically (Berglund et al., 2003) and influence ecological impact and disease susceptibility.

As ecosystem engineers, macroalgae are susceptible to infectious diseases triggered by abiotic and/or biotic environmental stressors (A Abdul Malik et al., 2020), with potentially significant ecological consequences. However, while diseases in substrate-attached macroalgae are extensively documented, diseases in free-floating macroalgal-communities remain virtually unstudied – a critical knowledge gap since drift communities typically comprise five to ten genera with one or two dominant species. Disease outbreak in a single dominant species could rapidly spread through the community, negatively impacting habitat provision, nutrient cycling, and food web dynamics, with cascading coastal ecosystem effects (Ramus et al., 2022; Ravaglioli et al., 2021).

Macroalgal disease outbreaks result from interactions between environmental stressors and pathogenic microorganisms, with temperature serving as a primary driver (Campbell et al., 2011; Loureiro et al., 2017). Climate change intensifies disease severity by simultaneously compromising host defenses and promoting pathogenic shifts in associated microorganisms (Campbell et al., 2011; Burge et al., 2013). These temperature-mediated diseases have reached alarming prevalence (Harvell et al., 1999; 2009; Egan et al., 2014; Paix et al., 2021), and can cascade throughout entire food webs (Campbell et al., 2011). Bleaching, the most commonly observed macroalgal diseases, is characterized by the localized pigment loss, reduced photosynthetic capacity, compromised growth, and increased vulnerability to herbivores (Campbell et al., 2014; Kumar et al., 2016). Several studies link bleaching incidence to elevated seawater temperature, which destabilizes host-microbe interactions and promotes opportunistic pathogenicity (Wright et al., 2000; Zozaya-Valdés 2017; Campbell et al., 2011). Free-floating macroalgal communities may be particularly vulnerable to bleaching due to their higher surface water temperatures relative to subtidal or intertidal regions.

*Halymenia floresii* (Clemente) C.Agardh is a native red algal species along the Yucatan peninsula coast of Mexico, with significant economic and ecological importance due to its lambda-carrageenan and phycoerythrin production and its ability to reduce inorganic nutrients in the integrated multi-trophic aquaculture (IMTA) systems (Godínez-Ortega et al., 2008; Robledo and Freile-Pelegrín 2011). Drift *H.floresii* populations are commonly found along the Gulf of Mexico from the early spring to the mid-summer months in small patches (Pihl et al., 1996; Correia, 2021), where bleaching symptoms -particularly localized apical pigment loss – have been noted but remain poorly studied. Understanding bleaching disease in *H. floresii* requires a holobiont approach examining bacterial diversity and abundance in diseased versus healthy specimens.

Our previous research established a foundation for understanding *H. floresii* host-microbe interactions, identifying Quorum Sensing (QS) members among 31 axenic bacterial strains cultured from *H. floresii* surface, confirming Quorum Sensing Interference (QSI) activity in surface metabolites and identifying *Vibrion owensii* as a putative opportunistic pathogen capable of inducing bleaching (A Abdul Malik et al., 2020a, 2020b). Building on this foundation, the present study aims to assess the microbiome in free-floating *H. floresii* across physiological states, healthy (Hfh), bleached (Hfb), and degrading (Hfd) specimens using high-throughput 16S rRNA gene sequencing to characterize bacterial community structure, Scanning Electron Microscopy (SEM) to visualize microbial load and surface topography and microbial network analysis to investigate the bacterial associations reorganize during bleaching. This represents the first descriptive assessment of the microbiome dynamics associated with bleaching disease in a drift red macroalga, *H. floresii* and advances our understanding of host-microbe organization under environmental stress in free-floating macroalgal communities.

## 2. Materials and Methods

### 2.1 Algal material collection

With a rigorous sampling, the drift *H. floresii* were collected during winter (December 2019) along a 3-4 km stretch of coastline near the CINVESTAV Coastal Marine Station in Telchac, Yucatán, México (21.3419° N, 89.2636° W). Collections were performed early in the morning, following a visual survey conducted the previous evening that confirmed the absence of drift, ensuring that the thalli collected were freshly drifting and had experienced minimal environmental exposure prior to sampling. The sampling coastline was divided into approximately 0.5 km intervals, and 6 individual plants corresponding to distinct health states – healthy (Hfh), bleached (Hfb), and degrading (Hfd), each in replicates were collected at each collection point (total initial pool: ∼ 30 – 35 specimens). Each specimen was handled using sterile nitrile gloves and examined *in situ* for pigmentation, tissue integrity, texture, presence of epiphytes, and signs of mechanical damage. Following laboratory inspection, only the strongest and most representative specimens for each health category were selected for analysis, i.e. n = 6 individual plants. The rigorous screening narrowed the samples to three high-quality biological replicates in two sets per condition (health, bleached, degrading), ensuring that the microbiome profiles reflected health state rather incidental damage or environmental artifacts.

Immediately after collection, thalli were gently rinsed with ambient seawater to remove loosely attached particles and debris. Samples were transferred into sterile polypropylene bags, kept shaded and transported to the laboratory in an insulated cooler with ice packs within 1-2 hours of collection. In the laboratory, thalli were processed immediately under sterile conditions. Each specimen was gently rinsed with sterile filtered seawater (30 psu, autoclaved and filtered 0.5 microns) to remove residual particulates. Subsamples for SEM and molecular analyses were excised using sterile scalpel and only one health category was processed at a time to avoid any cross-contamination.

### 2.2 Scanning Electron Microscopy

In this study, we observed the topography of the *H. floresii* surface and enumerated the surface-associated microbiome on healthy, bleached and degrading *H. floresii.* For SEM we used small segments of *H. floresii* and rinsed thrice with 1X Phosphate Buffered Saline (pH – 7.2), to remove any loosely attached bacteria. The specimens were fixed in 2.5% glutaraldehyde in 0.1M sodium cacodylate buffer. The fixed specimens were washed five times with 1X PBS (pH – 7.2) and then twice with distilled water to completely remove the residues of glutaraldehyde. After dehydration in a gradient ethanol series (30%, 50%, 70%, 90%, and 100%) the samples were critical point dried with carbon dioxide at 31° C under 1100 psi and observed for any deformation under the microscope. After they proceeded to metallization of the samples with gold-palladium by sputter coating. The specimens are observed under a scanning electron microscope and pictures were taken. All the samples are analyzed as four replicates at five different angles. The SEM count data were analyzed by Regression to determine the significance of the results obtained.

### 2.3 DNA Extraction and 16S rRNA Gene Sequencing

DNA was extracted with the DNeasy PowerSoil (Qiagen, Hilden, Germany) commercial kit using 0.5 g of macroalga samples according to the manufacturer’s instructions. To obtain sufficient DNA for each biological replicate, 0.25 g of tissue from two individual plants was extracted separately and the resulting extracts were pooled to form one replicate. The quality of DNA extraction was assessed with 1% agarose gel. The amplification of the V3 and V4 hypervariable regions of the 16S rRNA gene was conducted using the bacterial primers set and the conditions suggested by Klindworth et al., (2013). Each PCR Reaction was performed in 20 µL final volumes, containing 10 µL of Phusion Flash High-Fidelity Master Mix (Thermo Scientific, Waltham, USA), 0.5 µL of each primer (10 μM), and 2 µL of extracted DNA (10–20 ng/μL).

PCR amplicons were purified with AMPure XP beads (Beckman Coulter Genomics, Brea, CA) and indexed using the Nextera XT kit v2 (Illumina, San Diego, CA, USA), according to Illumina’s 16S Metagenomic Sequencing Library Preparation protocol and then were quantified with a Qubit 3.0 fluorometer (Life Technologies, Carlsbad, CA, USA). The correct size of the amplicons was verified by capillary electrophoresis at QIAxcel Advanced (QIAGEN, Valencia, CA, USA). Individual amplicons were diluted on 10 mM Tris (pH 8.5) and pooled at equimolar concentrations (4 nM). Paired-end sequencing was performed in the CINVESTAV-Merida on the Illumina MiSeq platform (Illumina, San Diego, CA, USA) using a MiSeq Reagent Kit V2 (500 cycles). Retrieved raw reads are available in the NCBI Sequence Read Archive (BioProject PRJNA976364).

### 2.4 Bioinformatic analyses / 16S rRNA amplicon sequence analysis

Clean and demultiplexed reads had a minimum length of 250bp. Data were examined with the QIIME2 (2019.1) pipeline (Caporaso et al., 2010). Elimination of chimeras, error correction, and denoising, were carried out with the DADA2 package, obtaining the representative amplicon sequence variants (ASVs) (Callahan et al., 2016; 2017). Taxonomic assignment of ASVs was done with the V-SEARCH classifier (Rognes et al., 2016), using the SILVA reference database (v. 138.1). The MAFFT algorithm (Katoh and Standley, 2013) performed the alignment of representative sequences, filtered for non-conserved and gapped positions, building a phylogenetic tree with the fasttree2 plugin (Price et al., 2010). The lowest read count number (7,100) was used to normalize all sample data. The obtained ASVs were later analyzed and visualized with the phyloseq (McMurdie and Holmes, 2013), and ggplot2 (Wickham, 2009) libraries, in the R-studio environment (https://www.r-project.org/). Differences among samples were assessed with a Principal Coordinate Analysis (PCoA), estimating the weighted and unweighted UniFrac distance (Lozupone and Knight, 2005). Alpha diversity indices (observed ASVs, Shannon and Simpson) were calculated from analyzed samples (Anderson, 2001). Microbial association networks at the genus level were constructed to investigate the interactions between bacterial taxa associated with the different health states of *H. floresii* using the NetCoMi R package (Peschel et al., 2021).

## 3. Results

### 3.1 Scanning electron microscopy of Healthy, Bleached, and Degrading H. floresii

Scanning electron microscopy revealed distinct differences in surface microtopography and microbial load among healthy, bleached, and degrading *H. floresii* specimens (Fig. 1). The surface of healthy *H. floresii* revealed a thick sheath of Extracellular Polymeric Substance (EPS) covering the cortical cells whereas the bleached surface exhibited nearly complete deterioration of this outer sheath. Degrading specimens showed an intermediate condition with a shredded net-like EPS structure (Fig. S1).

**Fig. 1.**
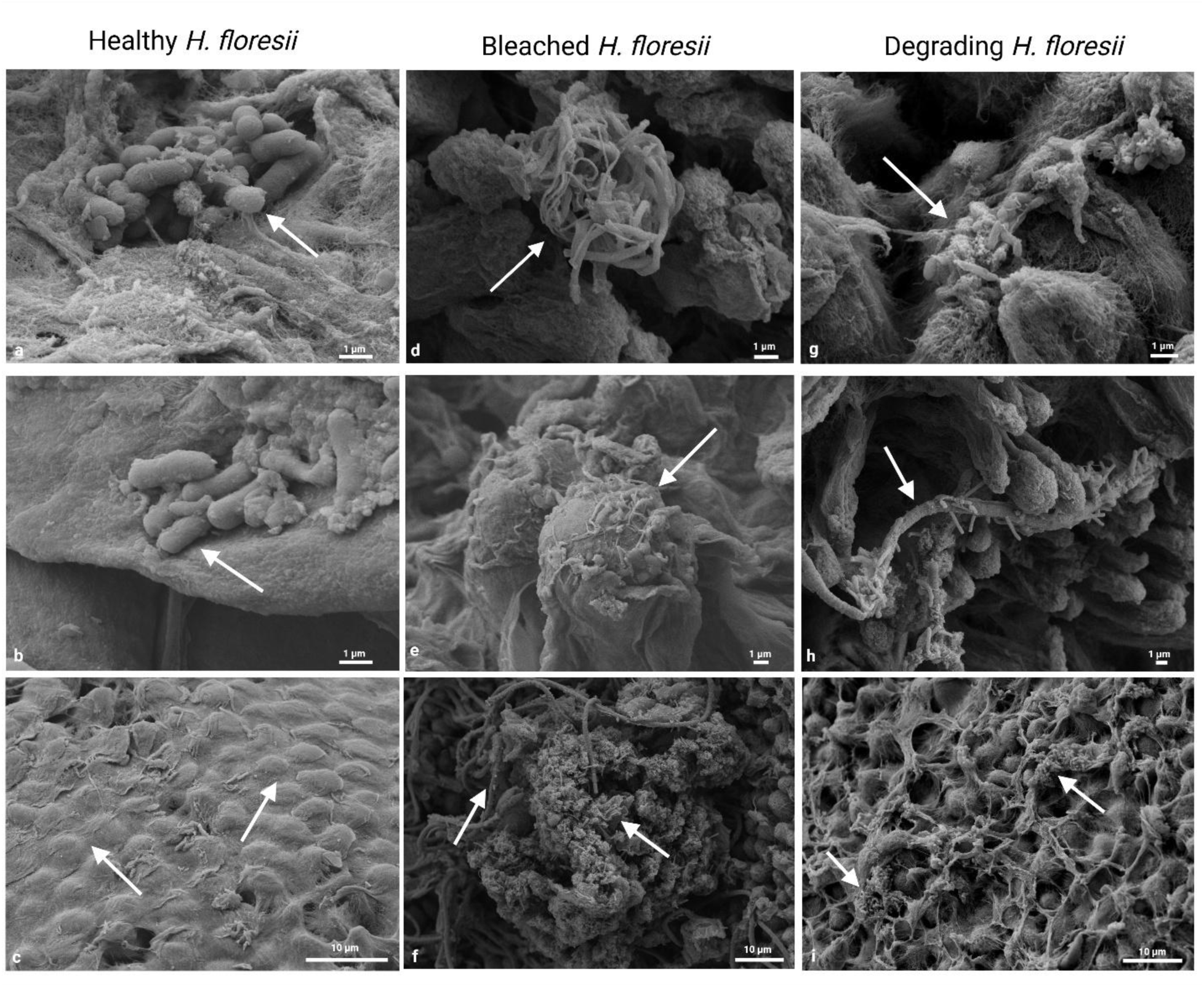
Scanning electron micrographs of reveal microbial density differences across health states of *Halymenia floresii*. Represenatative SEM images of (a,b.c) healthy, (d,e,f) degrading, and (g,h,i) bleached *H. floresii* thalli in triplicate samples. Images were at two magnifications: 1 μm and 10 μm. Healthy specimens (a-c) display intact extracellular polymeric substances (EPS) covering coritcal cells with low microbial density. Degrading specimens (d-f) exhibit shredded, net-like EPS deterioration with highest microbial colonization. Bleached specimens (g-h) show nearly complete EPS degradation with intermdediate microbial density. Scale bars: 1 μm and 10 μm as indicated. Microbial density ratios among health states (healthy:degrading:bleached) were 0.1:1.0:0.3 (R^2^ = 0.913, p<0.05).

The microbial density on the surfaces of healthy, bleached, and degrading *H. floresii* differed significantly, with observed ratios of 0.1:0.3:1.0, respectively. Statistical analyses of SEM counts confirmed significant differences among the three conditions (R^2^ = 0.913, p< 0.05) (Fig. S2). Notably, healthy *H. floresii* surfaces exhibited lowest microbial density while degrading showed the highest and bleached surfaces intermediate density.

### 3.2 Bacterial community structure analysis

High-throughput sequencing of the 16S rRNA gene yielded 122,506 bacterial raw reads. After quality filtering, including merging, denoising, and chimera removal using DADA2, a total of 111, 682 high-quality tags were retained for downstream analysis across healthy, bleached, and degrading samples. These sequences were clustered into amplicon sequence variants (ASVs) for taxonomic assignment.

Principal Component Analysis (PCoA) calculated on weighted and unweighted UniFrac distance matrix demonstrated clear separation of bacterial communities according to the health status of *H.floresii*, (Fig S3). The three sample types – healthy (Hfh), bleached (Hfb) and degrading (Hfd), – formed distinct clusters, indicating significant compositional differences in their associated microbiome. The weighted UniFrac analysis, which accounts for phylogenetic relatedness and relative abundance, showed stronger separation that unweighted analysis, indicating that both community membership and abundance structure differ among health status.

### 3.3 Bacterial community diversity across all health states of H. floresii

Taxonomic classification of bacterial ASVs revealed a total of 8 phyla, 10 classes, 15 orders, 20 families, and 23 genera across all *H. floresii* samples. The number of observed bacterial ASVs ranged from 18 to 194 (Table 1), with highest ASV richness detected in degrading specimens, intermittent richness in bleached samples, and lowest richness in healthy hosts (Fig. 2).

**Fig. 2.**
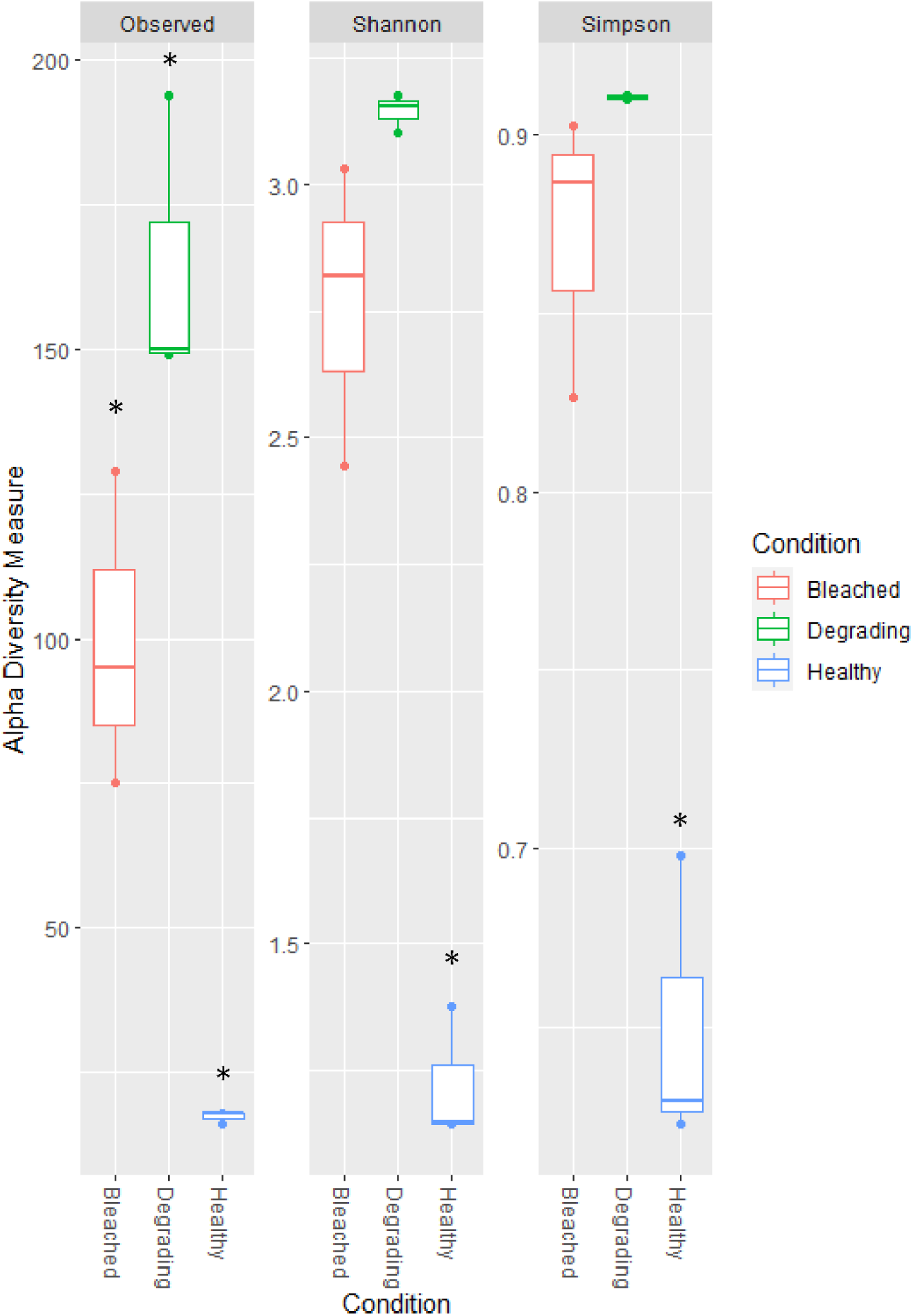
Alpha diversity indices calculated from retrieved 16s rRNA gene sequences. Shannon diversity index (middle panel) and Simpson diversity index (left panel) calculated from 16S rRNA gene amplicon sequences across healthy (Hfh, n = 3), degrading (Hfd, n = 3), and bleached (Hfb, n = 3) *H. floresii* samples. Box plots show median (center line), interquartile range (box), and range (whiskers). Shannon diversity increased significantly from healthy (1.3 – 1.4) to bleached (2.4 – 3.0) and degrading (3.1 – 3.2) samples. Simpson diversity indices ranged from 0.90 – 0.98, indicating increased community evenness in diseased states. Asterisks indicate statistically significant differences among health states.

**Table 1.**
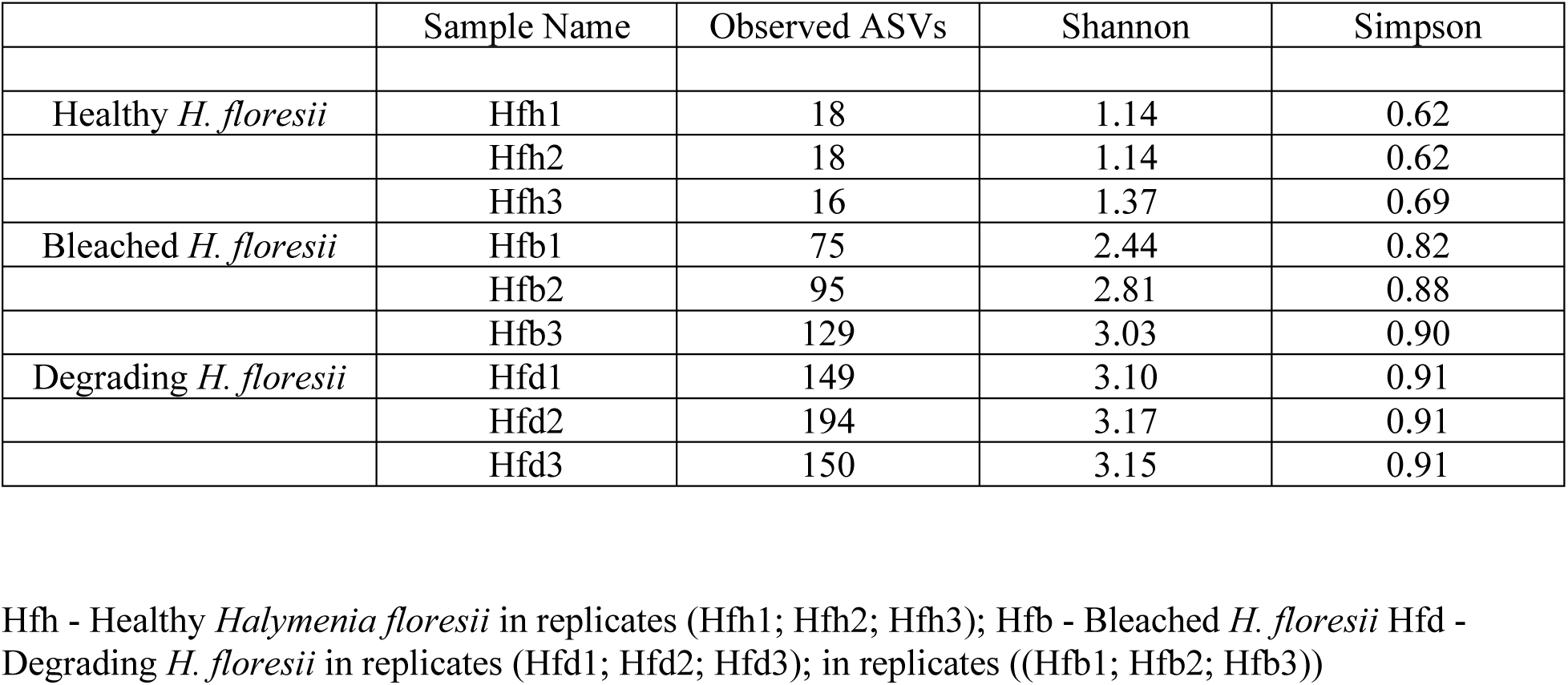
Observed ASVs; Shannon and Simpson indices of bacterial taxa associated with *Halymenia floresii* (n = 3)

Alpha diversity metrics varied across host health status (Fig. 2). Shannon diversity indices ranged from 1.1-1.3 in healthy samples, 2.4 – 3.0 in bleached samples and 3.1 – 3.15 in degrading samples. Simpson diversity indices ranged from 0.62 - 0.91 across all samples, with patterns parallelling Shannon diversity. These differences in alpha diversity were statistically significant among health status: degrading samples exhibited the highest diversity values, while healthy samples showed the lowest.

Bacterial community composition at the phylum level varied markedly across healthy, bleached, and degrading *H. floresii* (Fig. 3a). Healthy holobionts were dominated by Cyanobacteria (17-52%) and Pseudomonadota (formerly Proteobacteria) (47-81%). Degrading and bleached specimens harbored Bacteroidota (5-35%) and Pseudomonadota (41 −88%) becoming the predominant phyla (Fig. 3(a)). The abundance of the Cyanobacteria decreased dramatically, representing only 6-9% and 2 - 3% in degrading and bleached *H. floresii,* respectively. Several phyla occurred exclusively or predominantly in degrading and bleached samples. Desulfobacterota was observed in bleached and degrading samples with an abundance of 0.4% - 1% but was absent from healthy specimens. Camphylobacterota was observed in bleached (2 - 3%) and degrading (9 −10%) samples, and Firmicutes in bleached (0.3 - 1%) and degrading (2 - 3%) samples while neither phylum was detected in healthy specimens (Table S3).

**Fig. 3.**
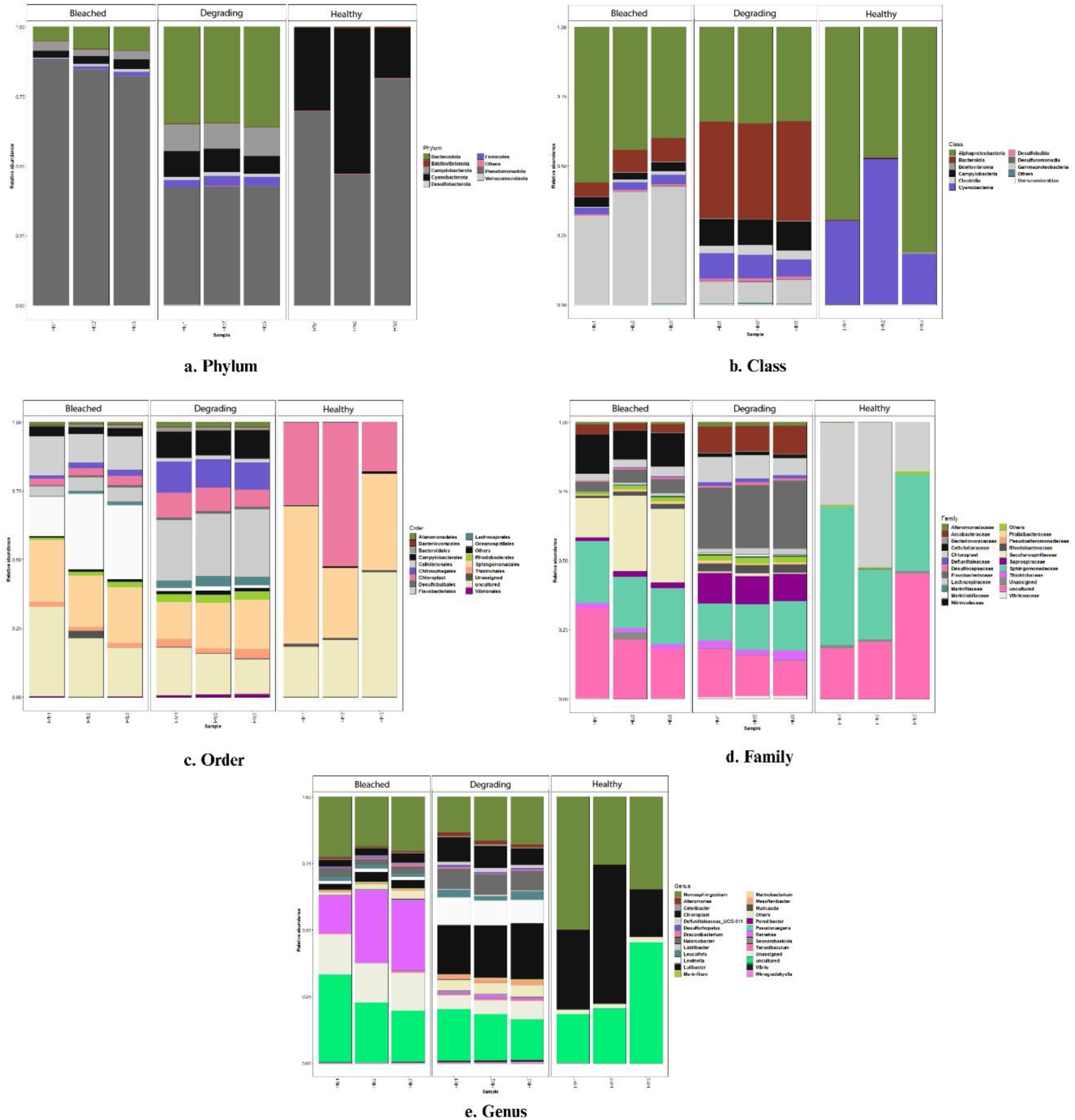
Bacterial community composition at multiple taxonomic levels. Stacked bar charts showing relative abundance of bacterial taxa at (a) phylum, (b) class, (c) order, (d) family, and (e) genus levels across bleached (Hfb1-3), degrading (Hfd1-3), and healthy (Hfh1-3) *H. floresii* specimens. Each bar represents one biological replicate. Healthy samples are dominated by Cyanobacteria (17 – 52%), Pseudomonadota (formerly, Proteobacteria) (41-81%, primarily Alphaproteobacteria from Sphingomonadales), Sphingomonadaceae (37%), and *Novosphingobium* (25 – 49%). Bleached samples show minimal Cyanobacteria (2-3%), elevated Gammaproteobacteria (8-38%), particularly Oceanospirillales), high Saccharospirillaceae (23%), and emergence of *Reinekea* (22.5%) with reduced *Novospingobium* (13-22%). Degrading samples exhibit loss of Cyanobacteria (6-9%), elevated Bacteroidota (up to 35%, primarily Bacteroidia from Flavobacteriales), emergence of Firmicutes (2-3%) and Campilobacterota (9-10%), dominance of Flavobacteraceae (23%), and high abundance of *Lutibacter* (20%). Overall, *Novosphingobium* enriched healthy communities, *Lutibacter* enriched degrading communities and *Reinekea* enriched bleached communities are observed here. Taxa <1% grouped as “Others”.

In the healthy *H. floresii* members of Bacteroidota (formerly known as Bacteroidetes) were not abundant (0.2%) but significantly increased in the bleached (8.4%) and degrading (35%) conditions. At the class level (Fig. 3b), healthy *H. floresii* primarily hosted Alphaproteobacteria (65%), comprising uncultured Alphaproteobacteria (28%) and members of the order Sphingomonadales (36%), along with Cyanobacteria (33.3%). In bleached and degrading hosts. Substantial change in the bacterial community structure is observed. Alphaproteobacteria remained present (34 – 46%) but with altered compositions, including uncultured Alphaproteobacteria (14.9 - 23.8%) and Sphingomonadales (15.9 −20.5%). Notably, Bacteroidia (7.1% to 35%) (from the orders Flavobacteriales (23.1-4.4%) and Chitinophagales (10.5-1.8%)), and Gammaproteobacteria (7.8 to 38%) (from the orders of Cellvibrionales (12.2-1.3%), Oceanospirillales (22.8-1.2%), Thiotrichales (2.6-1.5%) and Vibrionales (0.8-0.1%)) emerged as major components in diseased samples. Degrading samples additionally exhibited members of Clostridia (3%) specifically from the order Lachnospirales (3%,), which was less abundant in bleached hosts (0.8%) and absent from healthy specimens (Table S4).

Bacterial ASVs from all three health conditions were assigned to 20 families. In healthy *H. floresii*, the family Sphingomonadaceae dominated (36.7%), represented primarily by the genus *Novosphingobium* (36%). While in degrading samples Flavobacteriaceae (23%) was the predominant family including the genera *Lutibacter* (19.8%), *Mesoflavibacter* (1.4%), and *Tenacibaculum* (0.6%). In bleached samples, the family Saccharospirillaceae (22.6%) became highly abundant, represented almost exclusively by the genus *Reinekea* (22.5%). Several families appeared exclusively in degrading and bleached samples including Rhodobacteraceae (1.3 – 2.9%), comprising unassigned Rhodobacteraceae (0.8 – 1.8%), *Celeibacter* (0.2 – 0.4%), and *Pseudoruegaria* (0.1 – 0.2%), as well as Arcobacteraceae (2.9 – 9.5%), represented by the genus *Halarcobacter* (2.4 – 7.5%) (Table S5).

At the genus level (Fig. 3e), *Novosphingobium* (25-49%) was abundant in healthy *H. floresii* specimens, while showing lower representation in degrading (13-17%) samples and bleached (18-22%). Gammaproteobacterial *Reinekea* was abundant (22.5%) exclusively in bleached samples and completely absent from healthy hosts. Across all health states, a substantial proportion of sequences belonged to uncultured members of Pseudomonadota, accounting for 18 - 45% in healthy, 17 - 32% in bleached samples, and 12 - 17% in degrading (Table S6).

### 3.4 Network anlaysis revealing microbial association

Network analysis using NetCoMi identified three distinct clusters of bacterial genera were identified within the network, represented by purple, green, and orange nodes. The purple cluster was primarily composed of *Unassigned Cyanobacteria*, *Novosphingobium*, *Peredibacter*, *Luteolibacter*, and *Winogradskyella*, taxa predominantly associated with healthy samples. In contrast, two clusters were enriched in degrading and bleached samples. The orange cluster included genera such as *Reinekea*, *Colwellia*, *Alteromonas*, and *Leucothrix*, while the green cluster comprised *Vibrio*, *Marinomonas*, and *Marinifilum*. Genera within the orange and green clusters were mainly detected in degrading and bleached states. These disease-associated clusters exhibited distinct network properties, characterized by weaker positive associations and increased network fragmentation whereas the healthy-associated cluster showed predominantly positive associations among member taxa. These patterns indicate a reorganization of microbial associations from tightly connected, cooperative networks in healthy hosts to more fragmented, opportunistic networks in diseased hosts indicates a fundamental shift in community structure linked to the health status of *H. floresii* (Fig. 4).

**Fig. 4.**
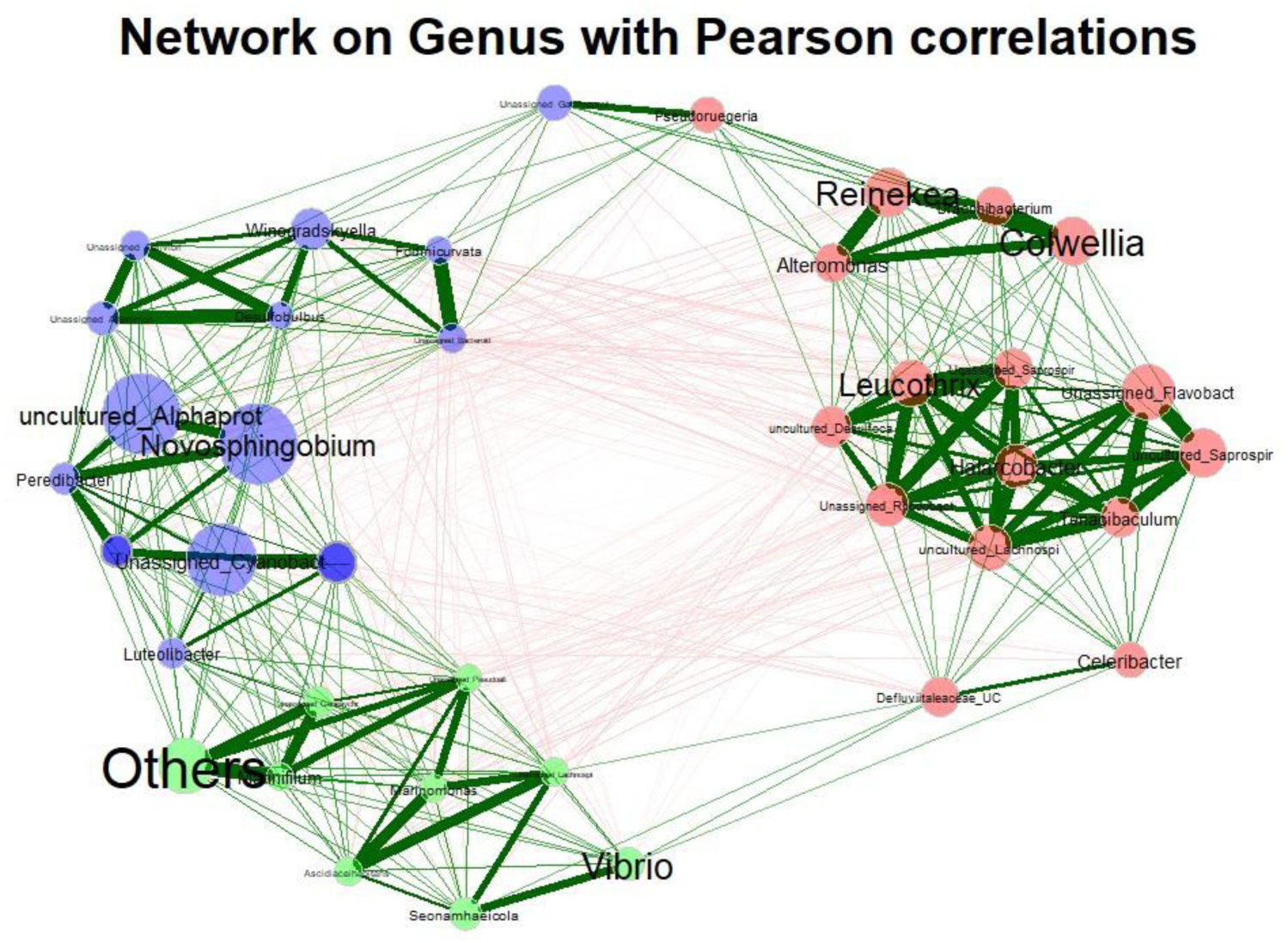
Microbial association network analysis reveals health-dependent reorganization of bacterial consortia. Network visualization of bacterial genus-level associations in *H.floresii* microbiomes constructed using NetCoMi analysis. Nodes represent bacterial genera, with node size proportional to relative abundance. Edge width indicates strength of association between genera. Three distinct clusters are identified: purple cluster (healthy-associated) dominated by uncultured Cyanobacteria, *Novosphingobium*, *Peredibacter*, *Luteolibacter*, and *Winogradskyella* with predominantly positive associations; orange cluster (disease-associated) including *Reinekea*, *Colwellia*, *Alteromonas*, and *Leucothrix*; green-cluster (disease-associated) comprising *Vibrio*, *Marinomonas*, and *Marinifilum*. The network topology demonstrates transition from tightly connected cooperative interactions in healthy states to fragmented, opportunistic networks in diseased states, supporting the pathobiome concept.

## 4. Discussion

Drift macroalgae originate from hard substrates like rocky shores but are transported by water activity and wave currents (Biber, 2007), introducing spatial and temporal heterogeneity into surrounding ecosystem. The ecological effects of drifting algal mats depend largely on their residence time in a specific location; however, a number of physical, chemical and biological factors may have bearing on their persistence (Berglund et al., 2003; Biber, 2007 & Arroyo and Bonsdroff, 2016). In this work, we focus on the most important biological factor, the associated microbiome of the algal host. This study provides the first descriptive assessment of microbiome transitions associated with bleaching disease in the drift red alga *H. floresii*. By integrating SEM, 16S rRNA sequencing, and microbial network analysis, the work characterizes how the structure and interactions of the algal-associated microbiome reorganize across healthy, bleached, and degrading thalli. The robustness of our microbiome assessment is underpinned by a highly rigorous and stringent sampling strategy for fresh drift material. DNA extractions were performed on tissue from two independent individuals and pooled to form each biological replicate, and triplicate replicates were generated for healthy, bleached and degrading thalli. Thus, minimizing bias from single-thallus variability and enhances the statistical reliability and reproducibility of the observed community patterns. Our findings demonstrate that drift *H. floresii* undergoes substantial microbiome restructuring during bleaching and degradation, marked by increased microbial diversity, loss of beneficial taxa, and the emergence of opportunistic bacteria, consistent with a pathobiome framework.

### 4.1 Scanning Electron Microscopy reveals the surface physiology

SEM observations (Fig. S3) showed that the macroalgal matrix is significantly deteriorated as a consequence of bleaching (see Supplementary Section). Extracellular polymeric substances (EPS) provide a matrix for primary attachment (Donlan, 2001), and the cell integrity at the surface determines the degree of microbial colonization. Surface roughness and microtopographical features function as selective determinants for microbial settlement and establishment, serving as mechanical antifouling defenses (Bers and Wahl 2004; Kientz et al., 2011; Wahl et al., 2012). Given these structural differences, it is noteworthy that healthy *H. floresii* surfaces exhibited lower microbial densities (0.1 relative ratio), while degrading surfaces showed the highest densities (1.0), and bleached surfaces intermediate (0.3), despite possessing a structurally intact EPS layer that might be expected to support colonization.

This inverse relationship suggests that healthy *H. floresii* may maintain active chemical defenses that complement its physical surface properties. Our previous metabolomic studies demonstrated that surface metabolites from *H. floresii* effectively quench quorum-sensing active molecules from bacterial isolates derived from this alga, indicating that the host actively suppresses microbial proliferation through interference with bacterial communication systems (A Abdul Malik et al., 2020a, b). This chemical defense strategy - maintaining a ‘microbially controlled’ surface – prevents biofilm formation and protects against microfouling while presumably allowing selective colonization by beneficial symbionts. Such spatial segregation of microbial communities has been documented in other macroalgae; for example, in green alga *Ulva reticulata* maintains lower bacterial fouling on healthy regions through waterborne polar molecules with antifoulant properties (Sergey and Qian, 2002; Harder et al., 2004; Dahms and Dobretsov 2017), and spatial variation in brown alga *Fucus vesiculosus* shows reduced biofilm thickness at apical tips compared to older thallus regions (Parrot et al., 2019). Our observations extend these findings to drift red algae and underscore the dynamic spatial distribution of bacterial partners across healthy and diseased thalli.

### 4.2 Phylogenetic differences and host specificity in the free-floating healthy, bleached and degrading H. floresii

High-throughput 16S rRNA sequencing revealed consistent and significant differences in bacterial community composition of free-floating healthy, bleached and degrading *H. floresii,* aligning with prior studies on substrate-attached diseased macroalgae. In red alga *Delisea pulchra*, bleaching-associated phylogenetic shifts primarily involved taxa from *Colwelliaceae, Rhodobacteraceae, Thalassomonas,* and *Parvularcula* (Fernandes et al., 2012). In contrast, *H. floresii* exhibited distinct family-level transitions centered on *Rubritaleaceae*, *Saccharospirillaceae*, *Saprospiraceae,* and *Arcobacteraceae*, suggesting that while bleaching processes may share common mechanisms across red algae, specific pathogenic consortia are host-dependent.

The observed distinction between phylogenetic diversity and apparent functional stability in macroalgal microbiomes arises from stochastic colonization by functionally equivalent but taxonomically distinct bacterial groups (Burke et al., 2011; Roth-Schulze et al., 2016; Korlević et al., 2021). This results in a two-tiered community structure: abundant core taxa are characterized by functional redundancy, whereas successional taxa are distinguished by host-specific functions (Korlević et al., 2021). In *H. floresii*, high-abundance taxa associated with healthy states – particularly *Novosphingobium* – likely possess functional traits that protect the host by carefully recruiting or competitively inhibiting the core members of the consortia. Conversely, successional taxa that dominate during disease progression are characterized by host-specific virulence mechanisms suited to exploit compromised algal physiology.

These observations support the emerging concept of pathobiome rather than single-pathogen causation. *Vibrio owensii* (MT176134) was previously identified as a significant bleaching-inducing pathogen in *H. floresii*; however, there were other strains (such as *Pseudoalteromonas arabiensis*, *P*. *mariniglutinosa*, *Ruegeria* sp., *Alteromonas* sp., *Epibacterium* sp., *Alteromonadaceae bacterium*, *Roseobacter* sp.) identified as potential pathogens (A Abdul Malik et al., 2022). Therefore, it might be time to shift the concept of “identified” pathogens to that of pathobiome in which the whole consortia contribute actively to the diseased condition. This perspective is further supported by recent work on *D. pulchra*, where bleaching resulted from synergistic interactions among opportunistic pathogens whose virulence traits are distributed across the consortium (Zozaya-Valdes et al., 2017).

### 4.3 Microbial network reorganization reflects health-dependent community dynamics

To further investigate these community-level dynamics beyond taxonomic shifts and relative abundance patterns, the microbial association network provides an additional ecological dimension by revealing how bacterial genera interact as part of a structured consortia rather than as isolated taxa. The NetCoMi-based network identified three distinct clusters of bacterial genera, which aligned closely with the health status of *H. floresii*. The cluster dominated by *Unassigned Cyanobacteria*, *Novosphingobium*, *Peredibacter*, *Luteolibacter*, and *Winogradskyella* was primarily associated with healthy samples, suggesting a tightly connected and potentially cooperative microbial assemblage. The prevalence of positive associations within this cluster supports the concept of functional redundancy and mutualistic interactions that stabilize the healthy holobiont (Bordenstein and Theis, 2015). In contrast, the clusters enriched in degrading and bleached samples, including genera such as *Reinekea*, *Colwellia*, *Alteromonas*, *Leucothrix*, *Vibrio*, *Marinomonas*, and *Marinifilum*, exhibited a reorganization of associations indicative of ecological destabilization. The emergence of these clusters reflects a shift from a cooperative network structure toward one dominated by opportunistic and potentially pathogenic taxa. This network-level reassembly provides strong support for the pathobiome concept discussed above, wherein disease is not driven by a single pathogen but by a consortium whose members collectively alter host–microbe interactions.

The abundance of *Novosphingobium* in bleached samples was significantly reduced compared to the healthy *H. floresii* (see Results section) and replaced by *Reinekea* (22.5%). The observed decline of *Novosphingobium* and its replacement by *Reinekea* in bleached samples gains additional relevance when viewed through the lens of microbial network structure. *Novosphingobium* occupies a central position in the healthy network, clustered with taxa primarily associated with positive correlations, consistent with its proposed roles in quorum sensing, surface defense, and microbiome stabilization (Wang et al., 2022). The loss of this genus from the network core during bleaching likely disrupts cooperative signaling and metabolic complementarities, reducing the resilience of the microbial community. Conversely, *Reinekea* occupies a distinct cluster associated with degrading and bleached states, pointing to its increased abundance is accompanied by a reconfiguration of microbial interactions rather than simple numerical dominance. Given its known capacity for polysaccharide degradation and opportunistic lifestyle (Avci et al., 2016), the network position of *Reinekea* supports the hypothesis that it contributes to host tissue breakdown not in isolation, but as part of a coordinated microbial assemblage. This shift from a symbiosis-dominated network to one structured around resource exploitation further underscores the dynamic nature of host–microbiome relationships under stress.

Comparable dynamics have been documented in other systems: seasonal studies of the brown alga *Sargassum thunbergii* observed higher *Reinekea* abundance under UV-B stress and lower *Novosphingobium* during summer months (Wang et al., 2023), while epiphytic communities on *Macrocystis pyrifera* show seasonal replacement of *Novosphinogbium* by *Ralstonia*, *Cyanobacterium,* and *Granulosicoccus* (Florez et al., 2019). Moreover, *Novosphingobium* strains were identified as QS bacteria with increased AHL production to protect the surface from xenobiotic compounds (Chen et al., 2020). These observations suggest that *Novosphingobium* may function as a microbial ‘second skin’, providing microbial defence to the host.

### 4.4 Pathogenicity on the surface of H. floresii’s

Gammaproteobacteria showed distinct abundance patterns across health states. Healthy specimens harbored predominantly Alphaproteobacteria with with negligible Gammaproteobacteria. In contrast, degrading and bleached samples contained substantial Gammaproteobacteria abundance, particularly from the orders of *Oceanospirillales* (*Reinekea*), *Vibrionales* (*Vibrio*), and *Cellvibrionales* (*Colwellia*, *Alteromonas*). Thus, Gammaproteobacteria emerging as key players in *H. floresii* bleaching pathogenesis. This parallels observations in *Eucheumatoid* species affected by Ice-Ice disease, where Gammaproteobacteria (*Pseudoalteromonas*, *Alteromonas, Stenotrophomonas*) and select Alphaproteobacteria (*Aurantimonas*, *Ochrobactrum*) comprise the pathogenic consortium (Ward et al., 2022), suggesting that certain Gammaproteobacteria may recurrently exploit stressed macroalgal hosts.

In addition, members of Bacteroidia (particularly *Tenacibaculum* and *Lutibacter*) appeared exclusively or predominantly in degrading and bleached samples, raising the possibility that they may function as secondary colonizers exploiting damaged tissue or contribute to further breakdown through enzymatic processing of algal polysaccharides (Hudson et al., 2022). In our previous work based on a culture-dependent study, we have also isolated and identified the same genera *Tenacibaculum* sp. (MT176141) and *Aquimarina* sp. (MT176152) from *H. floresii,* and both were confirmed as quorum sensing bacteria (A Abdul Malik et al., 2020a). This linkage between quorum sensing and pathogenicity - mechanism governing both symbiosis and virulence - suggests that dysregulation of bacterial cell-cell communication constitutes a critical transition point during disease development. Such complexity of bleaching disease in *H. floresii* – involving multiple bacterial phyla with distinct ecological roles – reinforces the pathobiome concept and suggest that effective disease management may require understanding not single pathogens but entire pathogenic assemblages.

### 4.5 Significance of Reinekea abundance in bleached samples and environmental drivers

Among the opportunistic taxa enriched in diseased samples, *Reinekea* possesses particular relevance to *H. floresii* bleaching given its documented capacity for algal polysaccharide degradation seasonality (Avci et al., 2017; Korlević et al., 2021). Members of this genus specialize in degrading mannan-type polysaccharide and carrageenan (Hakamada et al., 2014), the primary structural component of *H. floresii* cell walls. Their emergence as dominant taxa in bleached specimens suggest they may actively degrade the host’s carrageenan matrix, exacerbating tissue degradation beyond that caused by initial bleaching. This represents a critical mechanistic link between microbiome reorganization and visible disease symptoms.

The observation that *Reinekea* abundance was substantially reduced in degrading (0.5%) compared to bleached (22.5%) samples is noteworthy, implying *Reinekea* is particularly associated with the acute bleaching phase rather than the chronic degradation that follows. Once extensive tissue breakdown occurs and algal biomass becomes highly degraded, *Reinekea* may yield metabolic advantages to other scavenging bacteria, or competition for limiting resources may reduce its competitive advantage. Such temporal dynamics indicate that disease progression involves distinct ecological phases, each with characteristic microbial assemblages. Given that *H. floresii* produces lambda-carrageenan with commercial value and serves ecological functions in coastal environments, bleaching-induced polysaccharide degradation carries significant implications. If validated, the hypothesis that *Reinekea* and related taxa actively degrade carrageenan during bleaching suggests that diseased populations may lose not only photosynthetic capacity but also structural integrity – potentially affecting habitat provision and nutrient cycling in coastal drift communities (Avci et al., 2017).

Temperature is widely recognized as a primary driver linking climate change to macroalgal bleaching diseases. Elevated water temperatures can increase the physiological stress in macroalgal hosts while favoring more virulent or opportunistic phenotypes is associated bacteria, a mechanism analogous to that described in many coral bleaching syndromes (Gleason & Wellington, 1993; Hughes et al., 2007; Webster et al., 2011). Free-floating macroalgae may experience particularly pronounced thermal stress compared to substrate-attached populations, as surface waters inhabited by drift communities experience greater temperature fluctuations and higher absolute temperatures relative to subtidal or intertidal zones. The Gulf of Mexico, where this study was conducted, represents a tropical system where even modest temperature increase can exceed critical thermal thresholds for local macroalgal species. Notably, higher *Reinekea* abundance has been documented in *Sargassum thunbergii* under UV-B radiation stress (Wang et al., 2023) and *Sargassum fusiformis* under salinity extremes (Dai et al., 2022), implying environmental stressors broadly select this opportunistic genus. If temperature-induced bleaching in *H. floresii* is mediated by increased *Reinekea* abundance coupled with declining *Novosphingobium* – representing a loss of host-protective microbial defenses – this mechanism would explain the consistent correlation between elevated temperatures and bleaching frequency across macroalgal systems.

## 5. Conclusion

This study provides the first detailed characterization of microbiome patterns associated with bleaching disease in the drift *Halymenia floresii.* Our findings underscore the vulnerability of drift macroalgae to disease processes intensified by environmental stress. This study characterizes microbiome patterns with different health states in *H.floresii,* revealing distinct bacterial assemblages in healthy, bleached, and degrading specimens; however, special attention is required to understand the question of causalities. A key finding is that healthy *H.floresii* maintains a highly specific, low-diversity microbiome dominated by protective symbionts, and the diseased specimens exhibit elevated diversity and loss of specificity, consistent with dysbiosis (Li et al., 2021). This pattern underscores that microbiome health depends not simply on niche occupation but on precise genetic and metabolic complementarities between host and symbionts. Taken together, the integration of microbial network analysis with taxonomic, and microscopic observations provides a more holistic understanding of bleaching in *H. floresii*, which is critically essential for the coastal aquaculture sector since these drift diseased algae serve as vectors for disease emergence. This work provides a foundational reference for understanding disease ecology in *H. floresii* and highlights the need for future functional studies to elucidate the biochemical and environmental drivers of bleaching in drift macroalgal systems.

## 6. Declarations

### Authors’ contributions

S.A.A.M - wrote the main manuscript text; S.A.A.M; D.R and J.Q - designed the experimental protocol; S.A.A.M and A.M.G – performed the experiments; S.A.A.M and S.C - analyzed the data, prepared the figures; D.R and N.B - provided the fund; All authors reviewed the manuscript.

### Availability of data and materials

Raw sequencing data have been deposited in the NCBI Sequence Read Archive under BioProject accession number PRJNA976364. The authors confirm that the data supporting the findings of this study are available within the article and its supplementary materials.

